# The immune-opioid axis in prediabetes: prediction of prediabetes with insulin resistance by plasma interleukin-10 and Endomorphin-2 to kappa-opioid receptors ratio

**DOI:** 10.1101/2020.06.26.173120

**Authors:** Shatha Rouf Moustafa

**Affiliations:** Clinical Analysis Department, College of Pharmacy, Hawler Medical University

**Keywords:** Prediabetes, insulin resistance, interleukin, endorphin, endogenous opioid receptor

## Abstract

**Background:** Prediabetes is characterized by a hemoglobin A1c of 5.7%–6.4% and fasting blood glucose of 100–125 mg/dl. A high percentage of prediabetes subjects develops into type 2 diabetes mellitus in the following years. The effect of opioid peptides and their receptors, in addition to immunological cytokines on prediabetes, is not well understood.

**Objective:** We hypothesize that opioid peptides and their receptors affect the insulin and the insulin resistance (IR) in patients with prediabetes and that the immune cytokines, IL-6 (inflammatory factor) and IL-10 (anti-inflammatory factor), influence the opioid system.

**Methods:** A total of 60 patients with prediabetes and IR (prediabetes+IR), 60 patients with prediabetes without IR (prediabetes-IR), and 60 controls participated in the study. The IR state was HOMAIR > 2.5. The enzyme linked immunosorbent assay was used to measure interleukin (IL)-6, IL-10, μ- and κ-opioid receptors (MOR and KOR), endomorphin-2 (EM2), and β- endorphin (βEP).

**Results:** The subjects with prediabetes had dyslipidemia, and not all of them underwent the IR state. The IL-6, IL-10, β-endorphin, MOR, and endomorphin-2 were higher in the prediabetes subgroups compared with the control group. MOR was correlated with IL-10 and KOR. Prediabetes+IR can be predicted by the increased levels of the combination of IL-10, βEP, and EM2 and by the combination of IL-10 and endomorphin-2/KOR with good sensitivity and specificity.

**Conclusion:** Opioid peptides and their receptors were upregulated in patients with prediabetes depending on the significance of IR. These changes in the opioid system depend on the immune cytokines. Both systems need to be normalized to prevent further development into diabetes mellitus.

## Introduction

Impaired fasting blood glucose (FBG) or glucose tolerance occurs years before developing into a clear type 2 diabetes mellitus (T2DM), and this state is known as a prediabetes, which is a considerable risk factor for developing diabetes [1]. More than one-third of American adults above the age of 20 years develop prediabetes, a condition in which the blood glucose or the hemoglobin A1c (HbA1c) and FBG levels are higher than normal but not sufficiently elevated to be grouped as diabetes [2,3]. In the most recent US guidelines, prediabetes is defined as HbAic of 5.7%-6.4% and FBG of 100-125 mg/dl [4]. Approximately 87 million American adults have prediabetes, and approximately 30%-50% develop T2DM [5]. One of the dangerous factors of prediabetes is insulin resistance (IR), which refers to the reduced sensitivity or reactivity of tissues to insulin-mediated biologic activity [6]. IR leads to high blood sugar levels, prediabetes, and T2DM. IR, impaired insulin action, and insulin hypersecretion are central to the pathophysiology of prediabetes [7]. However, not all patients with prediabetes experience a sufficient increase in IR to be classified as an insulin-resistant patient. The clear condition of IR is when the HOMAIR value is more than the cutoff value (> 2.5) [8,9]. As the blood glucose increases, it joins the hemoglobin and elevates the level of HbA1c. This glycated hemoglobin is associated with the roles of IR [10].

Many evidence associates insulin with neuron functions and structure. Insulin signaling is important for neuronal survival, learning, and memorization. In animal models, impaired insulin signaling leads to a collection or gathering of Aβ oligomers [11,12]. Upon formation, Aβ oligomers exhaust insulin receptors from the surface membrane of neurons, thereby contributing to IR [13]. and generating an abnormal phosphorylation of insulin receptor substrate (IRS). This condition reduces the normal prosurvival signaling in neurons and promotes their death by apoptosis [14]. Notably, most of these studies focus on brain IR rather than systemic IR. These structural network abnormalities are related to the delay of the information processing speed [15]. IR occurring during midlife may elevate the danger of cognitive decline later in life. Elevated IR is associated with increased risk of cognitive decline, including impaired verbal fluency and slow simple reaction time [16]. Therefore, the relationship between IR roles and the molecules that have a function in the brain duties in some insulin-related disorders, including prediabetes, remains to be elucidated. These groups of molecules include endogenous peptides (e.g., β-endorphin and endomorphin-2) and their receptors, μ- (MOR) and κ-opioid (KOR) receptors. Another significant subject is the immune system molecules especially interleukin (IL)-6 and IL-10. In addition to yielding analgesia, glucose homeostasis is regulated by opioids through changing the insulin secretion [17]. The demonstration of β-endorphins in the human endocrine pancreas [18], and the possibility of causing insulin and glucagon liberation in animals and humans [19], suggest that opioids may affect glycoregulation [17]. Opioid receptors are expressed in the pancreas and on the pancreatic alpha and beta cells [20], thereby affecting endogenous opioid-mediated glucose and insulin homeostasis [21, 22]. An interaction occurs between the MOR and the IRS 1 in muscle tissues and cells [23]. Moreover, disorders related to the IR line polycystic ovary syndrome are accompanied with increased plasma immunoreactive β-endorphins [24]. On the basis of the linear regression analysis, the strongest proteins related to beta-cell function/HOMA-IR is the beta-endorphin [25]. The beta-endorphin is an influential variable in the ROC curve analysis for discrimination between high and low beta-cell functions and has a strong relationship with the beta-cell function, as observed in regression analysis [25]. Therefore, endogenous opiates are partially responsible for hyperinsulinemia and IR in prediabetes. The analysis of the isolated islets indicates that islet insulin hypersecretion is mediated directly by MOR, which is expressed on islet cells via a mechanism downstream of the ATP-sensitive K channel activation by glucose. Therefore, MOR manages the body weight by insulin release [26]. Opioids influence the beta cell function, as observed in individuals using heroin (MOR agonist) with elevated blood glucose levels [27]. Increased glucose utilization and decreased hepatic gluconeogenesis following the enhancement of peripheral MOR and the variation of genes in glucose metabolism [28] are among the proposed mechanisms. The enhancement of the chronic activation of opioid receptors stimulates hyperglycemia [29]. Puerarin enhances MOR expression and phosphorylation and elevates insulin-stimulated the glucose transporter 4 (GLUT4) translocation to the plasma membrane in the skeletal muscle of diabetic rats [23]. This study first supports the interaction between the MOR and the IRS 1 in the muscle tissues and cells [23]. The mechanism of enhancing IR may be associated with rising cell-surface level of GLUT4 by decreased FBG and fasting serum insulin levels and increased GLUT4 trafficking and intracellular insulin signaling [30]. The EM-2-induced inhibition of the gastrointestinal transit in diabetic mice is significantly weakened compared with that in nondiabetic mice. The inhibitory effects of EM-2 are mediated by MOR, which exerts a reduced role in diabetes. Moreover, poor blood glucose control may result in the attenuated effects of EM-2 [31].

IL-6 was studied because the impairment of the insulin-degrading enzyme (IDE) is associated with obesity and T2DM. IL-6 and IDE concentrations are significantly increased in the plasma of humans after an acute exercise compared with the pre-exercise values. Positive correlations between IL-6 and IDE activity and between IL-6 and IDE protein expression are observed. Our outcomes denote the novel role of IL-6 on insulin metabolism, suggesting new potential therapeutic strategies and focusing on insulin degradation, for the treatment and/or prevention of diseases linked to hyperinsulinemia, such as obesity and T2DM [32].

Interleukin-6 (IL-6) is a proinflammatory cytokine that conclusively motivates the progression of IR and the pathogenesis of T2DM through the generation of inflammation by controlling differentiation, migration, proliferation, and cell apoptosis [33]. The presence of IL-6 in tissues is a normal consequence, but its irregular generation and long-term exposure leads to the development of inflammation, which stimulates IR and overt T2DM [33]. A mechanistic association occurs between the promotion of IL-6 and IR. IL-6 is the purpose of IR by impairing the phosphorylation of insulin receptor and IRS-1 through the stimulation of the expression of SOCS-3, a potential inhibitor of insulin signaling [33]. IL-10 is a cytokine with antiinflammatory characteristics that can modulate inflammatory responses by repressing the generation of proinflammatory cytokines [34]. Serum IL-10 has a converse association with hyperinsulinemia and IR as IL-10 decreases with an increase in HOMA-IR [35].

This study was performed because the direct link between opioid and opioid receptors in patients with prediabetes is still weak, and molecular, physiological, and clinical studies are needed to determine the role of the opioid system in the etiology and pathogenesis of prediabetes.

This clinical study was designed to add clinical information on the diagnosis, prevention, and identification of the risk factors related to the initial occurrence of prediabetes to improve the quality of life by enhancing the health care and education of individuals. The results of the current study may make a difference in the care of future individuals. The association of these factors with prediabetes remains to be elucidated. We hypothesized that opioid peptides and their receptors affect the insulin and IR functions of patients with prediabetes. Moreover, we explored the effects of immune cytokines IL-6 (inflammatory factor) and IL-10 (anti-inflammatory factor) on the opioid system and IR parameters in patients with prediabetes.

The MOR is hypothesized to have an important role in IR. Several reports have investigated the role of MOR in insulin sensitivity and IR, but the results are somehow controversial. Thus, this study was conducted. This research is essential for the pharmacological intervention for the potential targeting of MOR for IR therapy.

## Materials and Methods

### Subjects

Patients: More than 500 subjects, who check FPG routinely in the laboratories, were examined to select our study group with restricted criteria. A total of 120 subjects with prediabetes were chosen to participate in the study. These subjects were diagnosed by a physician in accordance with the American Diabetes Association’s criteria (FPG = 5.55–6.94 mM, HbA1c = 5.7%–6.4%). The samples were collected from private clinics and laboratories and the Rizgari Teaching Hospital in Erbil City, Kurdistan Region, Iraq from October 2019 to December 2019. The patients fasted for 8–12 h. All procedures were conducted in accordance with the established ethical standards. All study subjects provided written informed consent prior to participation in the study. The study was carried out in accordance with the international and Iraq ethics and privacy laws and approved by the Ethics Committee of Medical Research at College of Pharmacy/Hawler Medical University. The study was performed in accordance with the ethical standards in the Helsinki Declaration of 1975, as revised in 2008, for experiments involving humans.

The subjects with prediabetes were further divided into two subgroups in accordance with the results of HOMA2IR. The first group, prediabetes+IR, comprised subjects with high IR state (HOMA2IR > 2.5). The second group, prediabetes-IR, comprised subjects with low IR state (HOMA2IR < 2.5). This classification occurred deliberately in the same number of subjects in the two groups to remove the possible bias of the number of cases. A total of 55 healthy adult subjects were selected as the control group. Age ranges and sex ratios were comparable to that of the prediabetes groups. None of these subjects manifested an evident systemic disease or took drugs. Furthermore, in all subjects, the C-reactive protein (CRP) was lower than 6 mg/l, which excluded overt inflammation. Tobacco use disorder was examined in accordance with the DSM-IV-TR criteria. Body mass index (BMI) was calculated using the formula: body weight (kg)/squared length (m^2^). Subjects performing more than 30 min of moderate activity for 2–3 times/week and never or less than one time per week were considered to do physical activity [4].

Exclusion criteria: The present study excluded patients who satisfied the following criteria: patients with serum TG ≥ 5.32 mM based on the Friedewald’s formula, FPG > 25 mM, and fasting insulin > 400 pM based on the HOMA calculator software requirements. Patients with evident major diabetic complications, such as heart diseases, and those receiving lipid-lowering medications (e.g., simvastatin or atorvastatin) and metformin were also excluded. Subject with abnormal blood pressure were also excluded. No subject had a urinary albumin/creatinine ratio of more than 30 mg/g to exclude microalbuminuria, which indicated microvessel damage.

### Measurements

All blood samples were collected in the morning after 12 h of fasting. After collection and separation, the serum samples were separated into three aliquots and stored in a refrigerator until use. The serum glucose, total cholesterol, and TG were determined spectrophotometrically by observing the enzymatic reactions by using commercially available kits (Spinreact^®^, Spain). Serum HDLc was determined after the other lipoproteins were precipitated using a reagent containing sodium phosphotungstate and magnesium chloride. The cholesterol contents in the supernatant were determined using a cholesterol kit. VLDLc was calculated on the basis of TG/2.19 and LDLc by using Friedewald’s equation (LDLc = Tc - HDLc - VLDLc). The HbA1c percentage in the whole blood and the urinary albumin/creatinine ratio were determined using the immunofluorescence analyzer *Finecare™* II *FIA* Meter (Guangzhou Wondfo Biotech Co., Ltd, China). The normal range of the HbA1c kit was 4%–6% and that of microalbumin ratio was ≤ 30 mg/g. The IR parameters were calculated on the basis of fasting glucose and insulin concentrations by using the HOMA calculator software (http://www.dtu.ox.ac.uk/homa-calculator/download.php). This software was used to generate the IR (HOMA2IR), insulin sensitivity (HOMA%S), and beta-cell function (HOMA%B) indices. An ideal normal-weight individual aged < 35 years had HOMA2IR of 1 and HOMA%B cell function of 100%. The serum insulin level was assayed using the insulin enzyme-linked immunosorbent assay (ELISA) kit, which is a solid-phase ELISA based on the sandwich principle, supplied by Calbiotech^®^, China. The serum levels of IL-10 (Elabscience^®^, Inc. CA, USA), MOR and KOR, endomorphin-2 (Mybiosource^®^, Inc. CA, USA), and IL-6 and β-endorphin (Melsin Medical Co, Jilin, China) were assayed using commercial ELISA kits. The sensitivities of β-endorphins, MOR, KOR, endomorphin-2, and IL-6 ELISA kits were 0.1, 7.18, 1.0, 0.33, and 0.1 pg/ml, respectively. All measured concentrations were greater than their assay sensitivities. All intra-assay coefficients of variation were < 10.0%. Serum CRP was measured using the kit supplied by Spinreact^®^, Spain through a test based on the latex agglutination principle.

### Statistical Analysis

Analysis of variance (ANOVA) was employed to assess the differences in all measured biomarkers between diagnostic categories, and the *χ*^2^ test was used to compare the proportions and nominal variables. The associations among variables were computed using the Pearson’s product-moment and the Spearman’s rank-order correlation coefficient. The multivariate general linear model (GLM) analysis was used to delineate the effects of diagnosis for the prediabetes+IR, prediabetes-IR, and control groups while controlling the background variables, including age and sex. Protected LSD tests were used to check pairwise comparisons among treatment means. The model-generated estimated marginal mean (SE) values were computed after adjusting for covariates. Multiple regression analysis was used to delineate the significant biomarkers associated with the prediabetes+IR, and the results were checked for multicollinearity (tolerance and VIF values) and homoscedasticity (White and Breusch-Pagan tests). The binary logistic regression analysis was used to delineate the important explanatory variables that predict prediabetes (versus control as reference group). The data were subjected to ln transformation to normalize the data distribution of the measured biomarkers (tested using the Kolmogorov-Smirnov test). However, the nonlinearity of the mean and variance of any biomarker is a predictable source of variability that is eliminated using *z* scores. The natural logarithm of the relevant *z* unit scores was computed to transform the nonparametric variables into normally distributed components and apply the statistical analysis as a linear group. All tests were two-tailed, and a p value < 0.05 was used for statistical significance. All statistical analyses were performed using the IBM SPSS windows version 25, 2017. Odds ratios (OR) and 95% confidence intervals (CI) for unfavorable glycemic status by study factors were calculated.

## Results

### Sociodemographic data

Table 1 shows the sociodemographic data of the prediabetes-IR, prediabetes+IR, and healthy control groups. No significant difference was observed among the groups in terms of age, BMI, sex ratio, employment, and marital status. The patients in the prediabetes-IR group were slightly more educated than those in the prediabetes+IR group.

**Table 1:** Demographic and clinical data of healthy controls (HC) and Prediabetes-IR versus Prediabetes+IR groups.

| Parameter | Control <sup>A</sup><br>N=58 | Prediabetes-IR <sup>B</sup><br>N=60 | Prediabetes+IR <sup>C</sup><br>N=60 | F/$\chi^2$ | df | p |
| --- | --- | --- | --- | --- | --- | --- |
| Sex (F/M) | 28/30 | 28/32 | 27/33 | 0.364 | 2 | 0.552 |
| Age year | 36.67±10.03 | 35.03±11.44 | 37.9±11.88 | 0.499 | 2/174 | 0.609 |
| BMI kg/m<sup>2</sup> | 26.39±3.72 | 27.76±4.01 | 27.63±5.14 | 0.909 | 2/174 | 0.407 |
| Smoking (Y/N) | 17/41 | 18/42 | 19/41 | 0.321 | 2 | 0.714 |
| Marital status (M/S) | 33/25 | 31/29 | 36/24 | 1.308 | 2 | 0.067 |
| Education years | 10.2 (4.7) | 10.3 (5.2) <sup>C</sup> | 9.9 (4.3) <sup>B</sup> | 2.107 | 2/174 | 0.054 |
| Employment (Y/N) | 36/22 | 40/20 | 38/22 | 1.163 | 2 | 0.088 |
| Physical activity (Y/N) | 21/37 | 20/40 | 18/42 | 1.412 | 2 | 0.056 |
| TC mM | 4.07±0.59<sup>C</sup> | 4.42±0.56 | 4.72±0.70<sup>A</sup> | 8.288 | 2/174 | 0.001 |
| TG mM | 1.40±0.14<sup>C</sup> | 1.49±0.17 | 1.57±0.36<sup>A</sup> | 3.701 | 2/174 | 0.029 |
| VLDLc mM | 0.64±0.07<sup>C</sup> | 0.68±0.08 | 0.72±0.16<sup>A</sup> | 3.701 | 2/174 | 0.029 |
| HDLc mM | 1.12±0.11<sup>C</sup> | 1.06±0.14 | 0.99±0.13<sup>A</sup> | 6.789 | 2/174 | 0.002 |
| LDLc mM | 2.32±0.60<sup>C</sup> | 2.69±0.58 | 3.01±0.74<sup>A</sup> | 8.725 | 2/174 | <0.001 |
| TC/HDLc | 3.69±0.64<sup>B,C</sup> | 4.27±0.81<sup>A,C</sup> | 4.81±0.80<sup>A,B</sup> | 16.512 | 2/174 | <0.001 |
| TG/HDLc | 1.27±0.19<sup>C</sup> | 1.44±0.26 | 1.61±0.42<sup>A</sup> | 9.298 | 2/174 | <0.001 |
| LDLc/HDLc | 2.11±0.60<sup>B,C</sup> | 2.61±0.75<sup>A,C</sup> | 3.07±0.81<sup>A,B</sup> | 13.214 | 2/174 | <0.001 |
| zLn IL-6 | -0.35±1.15<sup>B,C</sup> | 0.16±0.90<sup>A</sup> | 0.19±0.86<sup>A</sup> | 4.562 | 2/174 | 0.019 |
| IL-10 pg/ml | 7.9±2.27<sup>B,C</sup> | 11.46±4.52<sup>A,C</sup> | 14.08±8.19<sup>A,B</sup> | 9.362 | 2/174 | <0.001 |
| zLn βEP | -0.50±0.39<sup>B,C</sup> | 0.37±1.39<sup>A</sup> | 0.13±0.76<sup>A</sup> | 6.834 | 2/174 | 0.002 |
| zLn MOR | -0.34±1.15<sup>B,C</sup> | 0.19±0.96<sup>A</sup> | 0.15±0.81<sup>A</sup> | 4.087 | 2/174 | 0.024 |
| zLn (βEP/MOR) | -0.09±1.11 | 0.10±1.02 | 0.01±0.87 | 0.279 | 2/174 | 0.757 |
| zLn EM2 | -0.50±0.89<sup>B,C</sup> | 0.16±0.98<sup>A</sup> | 0.34±0.96<sup>A</sup> | 6.615 | 2/174 | 0.002 |
| zLn KOR | -0.35±1.20<sup>B,C</sup> | -0.21±1.14<sup>A</sup> | 0.14±0.64<sup>A</sup> | 3.964 | 2/174 | 0.031 |
| zLn (EM2/KOR) | -0.15±1.14 | -0.038±0.94 | 0.19±0.90 | 0.888 | 2/174 | 0.415 |
| HbA1c % | 5.01±0.43<sup>B,C</sup> | 6.02±0.22<sup>A</sup> | 6.17±0.2<sup>A</sup> | 132.746 | 2/174 | <0.001 |
| Glucose mM | 5±0.71<sup>B,C</sup> | 5.92±0.59<sup>A,C</sup> | 6.49±0.39<sup>A,B</sup> | 49.734 | 2/174 | <0.001 |
| Insulin pM | 68.54±10.79<sup>B,C</sup> | 100.74±13.64<sup>A,C</sup> | 140.48±12.38<sup>A,B</sup> | 256.628 | 2/174 | <0.001 |
| I/G mM | 14.07±3.40<sup>B,C</sup> | 17.12±2.46<sup>A,C</sup> | 21.79±2.84<sup>A,B</sup> | 52.921 | 2/174 | <0.001 |
| zLnHOMA2%B | 0.13±1.39 | 0.18±0.84 | 0.05±0.63 | 0.764 | 2/174 | 0.469 |
| HOMA2%S | 80.72±13.82<sup>B,C</sup> | 52.63±7.51<sup>A,C</sup> | 36.85±2.80<sup>A,B</sup> | 174.254 | 2/174 | <0.001 |
| HOMA2IR | 1.27±0.19<sup>B,C</sup> | 1.94±0.27<sup>A,C</sup> | 2.73±0.22<sup>A,B</sup> | 303.865 | 2/174 | <0.001 |

<sup>A,B,C</sup>: pairwise comparisons between group means.
zLn: z-score of the natural logarithm, FPG: fasting plasma glucose, TC: serum total cholesterol, TG: serum triglycerides, VLDLc: very low density lipoproteins, HDLc: high density lipoproteins, LDLc: Low density lipoproteins, IL: interleukin; KOR: κ-opioid receptor; MOR: $\mu$ opioid receptor; EM2: Endomorphin-2; βEP: β-endorphin; BMI: Body mass index, HOMA2IR: homeostasis model assessment 2 of insulin resistance, HOMA2%S: homeostasis model assessment 2 of insulin sensitivity percentage:
HOMA2%B: homeostasis model assessment 2 of beta-cell function percentage, IR: insulin resistance (HOMA2IR>2.5),

A significant increase (p < 0.05) in TC, TG, VLDLc, and LDLc and a significant decrease in HDLc were observed in the prediabetes+IR group compared with the control group. No such difference was observed between the prediabetes-IR and the prediabetes+IR groups. The atherogenic indices, TC/HDLc and LDLc/HDLc, were significantly different among the three study groups, and the scores followed the order: controls < prediabetes-IR < prediabetes+IR. The TG/HDLc showed a significant increase in prediabetes+IR group compared with the control group.

No significant difference was observed in the levels of the ratios of opioids to their receptors zLn (βEP/MOR) and zLn (EM2/KOR) among the study groups. The level of other opioids in the prediabetes groups was higher than that in the control group: zLn βEP (*F* = 6.834, df = 2/174, p = 0.002), zLnMOR (*F* = 4.087, df = 2/174, p = 0.024), zLnEM2 (*F* = 6.615, df = 2/174, p = 0.002), and zLnKOR (*F* = 3.964, df = 2/174, p = 0.031).

Serum IL-10 was significantly different among the three study groups (*F* = 9.362, df = 2/174, p < 0.001), and the score followed the order: controls < prediabetes-IR < prediabetes+IR. zLnIL-6 was significantly increased in the prediabetes groups compared with the control group, whereas no such difference was observed between the prediabetes subgroups.

The IR parameters, namely, glucose (*F* = 49.734, df = 2/174, p < 0.001), insulin (*F* = 256.628, df = 2/174, p < 0.001), I/G (*F* = 52.921, df = 2/174, p < 0.001), HOMA2%S (*F* = 174.254, df = 2/174, p < 0.001), and HOMA2IR (*F* = 303.865, df = 2/174, p < 0.001), followed the order: controls < prediabetes-IR < prediabetes+IR. The HbA1c% in the prediabetes-IR and the prediabetes+IR groups was higher than that in the control group (*F*=132.746, df = 2/174, p < 0.001). zLnHOMA2%B showed no significant difference among the study groups (*F* = 0.764, df = 2/174, p = 0.469).

### Differences in the biomarkers between the study groups

In the entire study group, significant correlations were observed between the zLnMOR and the following parameters: IL-10 *(r* = 0.306, p = 0.017), zLnKOR *(r* = 0.311, p = 0.016), and zLnIL-6 *(r* = 0.368, p = 0.004). zLnKOR was correlated with IL-6 (*r* = 0.451, p < 0.001) and zLnEM2 with (*r* = 0.377, p = 0.003).

### Multivariate GLM analysis

Table 2 displays the outcomes of the multivariate GLM analysis comparing the differences in the measured biomarkers among the three study groups while adjusting for age, BMI, sex, physical activity, education, IR parameters, and smoking. Significant differences (p = 0.040) were observed in the biomarkers among the groups with an effect size of 0.198, whereas the other covariates had no significant effects (p > 0.05). The tests for between-subject effects in Table 2 and the results in Table 3 showed the SE values and indicated that all eight biomarkers of the patients with prediabetes were significantly higher compared with those of the control group. Furthermore, the IL-0, zLnβEP, zLnMOR, and zLnEM2 in the prediabetes subgroups were significantly higher compared with those in the control group. Among all the examined biomarkers, zLnβEP had the highest effect on the diagnosis of prediabetes (*F* = 5.128, df = 2/174, p = 0.027, partial *η*^2^ = 0.067).

**Table 2:** Results of multivariate GLM analysis showing the associations between biomarkers and diagnosis while adjusting for background variables

| Type | Dependent variable | Explanatory variable | F | df | p | Partial $\eta^2$ |
| --- | --- | --- | --- | --- | --- | --- |
| Multivariate | IL-10, zLn βEP, zLn (βEP/MOR), zLn IL-6, zLnMOR, zLnEM2, zLnKOR, zLn (EM2/KOR) | Diagnosis | 2.203 | 16/320 | 0.045 | 0.192 |
|  |  | Sex | 0.450 | 8/168 | 0.867 | 0.046 |
|  |  | Age | 1.138 | 8/168 | 0.351 | 0.109 |
|  |  | BMI | 1.803 | 8/168 | 0.102 | 0.163 |
|  |  | Smoking | 0.604 | 8/168 | 0.750 | 0.061 |
|  |  | Marital status | 1.426 | 8/168 | 0.210 | 0.133 |
|  |  | Physical activity | 0.798 | 8/168 | 0.612 | 0.084 |
|  |  | HbA1c | 1.076 | 8/168 | 0.389 | 0.104 |
|  |  | Glucose | 0.801 | 8/168 | 0.590 | 0.079 |
|  |  | Insulin | 0.851 | 8/168 | 0.550 | 0.084 |
|  |  | I/G | 0.968 | 8/168 | 0.462 | 0.094 |
|  |  | HOMA2%S | 0.495 | 8/168 | 0.835 | 0.051 |
|  |  | HOMA2IR | 0.841 | 8/168 | 0.558 | 0.083 |
|  |  | zLnHOMA2%B | 1.177 | 8/168 | 0.328 | 0.112 |
|  |  | TC, TG, VLDL, HDL, TC/HDL, TG/HDL, and LDL/HDL | 0.002 | 8/168 | 0.966 | 0.001 |
| Tests of Between-Subjects Effects | IL-10 | Diagnosis | 2.102 | 2/174 | 0.152 | 0.029 |
| | zLn βEP | Diagnosis | 5.128 | 2/174 | 0.027 | 0.067 |
| | zLn (βEP/MOR) | Diagnosis | 0.884 | 2/174 | 0.350 | 0.012 |
|  | zLn IL-6 | Diagnosis | 0.202 | 2/174 | 0.654 | 0.003 |
|  | zLnMOR | Diagnosis | 1.131 | 2/174 | 0.291 | 0.016 |
|  | zLnEM2 | Diagnosis | 0.003 | 2/174 | 0.957 | 0.001 |
|  | zLnKOR | Diagnosis | 3.745 | 2/174 | 0.057 | 0.050 |
|  | zLn (EM2/KOR) | Diagnosis | 2.213 | 2/174 | 0.141 | 0.030 |

zLn: z-score of the natural logarithm, FPG: fasting plasma glucose, IL: interleukin; KOR: κ-opioid receptor; MOR: $\mu$ opioid receptor; EM2: Endomorphin-2; βEP: β-endorphin.

**Table 3:** Model-generated estimated marginal means values (SE) of the biomarkers in Prediabetes (versus healthy controls) and Prediabetes-IR versus Prediabetes+IR and healthy controls

| Biomarkers | Control <sup>A</sup> | Prediabetes-IR <sup>B</sup> | Prediabetes+IR <sup>C</sup> |
| --- | --- | --- | --- |
| IL-10 | 7.981(1.062)<sup>B,C</sup> | 11.461(1.026)<sup>A</sup> | 14.082(1.026)<sup>A</sup> |
| zLn βEP | 12.146(2.308)<sup>B,C</sup> | 23.652(2.230)<sup>A</sup> | 20.638(2.230)<sup>A</sup> |
| zLn (βEP/MOR) | 5.362(0.677) | 5.976(0.654) | 5.572(0.654) |
| zLn IL-6 | 0.314(-0.187) | 0.157(0.180) | 0.194(0.180) |
| zLnMOR | 0.431(-0.180)<sup>B,C</sup> | 0.194(0.174)<sup>A</sup> | 0.149(0.174)<sup>A</sup> |
| zLnEM2 | 0.518(-0.180)<sup>B,C</sup> | 0.163(0.174)<sup>A</sup> | 0.338(0.174)<sup>A</sup> |
| zLnKOR | 0.362(-0.186) | 0.213(0.180) | 0.140(0.180) |
| zLn EM2/KOR | 0.158(-0.191) | 0.038(-0.185) | 0.188(0.185) |

<sup>A,B,C</sup>: pairwise comparisons between group means
zLn: z-score of the natural logarithm, IL: interleukin; KOR: κ-opioid receptor; MOR: $\mu$ opioid receptor; EM2: Endomorphin-2; βEP: β-endorphin.

Table 4 shows the results of two binary logistic regression analyses examining the best predictors of prediabetes (versus controls) and prediabetes-IR (versus prediabetes+IR) by using an automatic stepwise method with biomarkers as explanatory variables while allowing the effects of other cofounders (age, sex, smoking, employment, and education). The first regression analysis showed that prediabetes was best predicted by increased levels of IL-10, zLnβEP, and zLnEM2 *(χ^2^* = 38.122, df = 7, p < 0.001, Nagelkerke = 0.480) with an accuracy of 76.7%, sensitivity of 85.0%, and specificity of 75.6%. The second regression analysis showed that the combination of IL-10 and zLnEM2/KOR were the best predictors of prediabetes+IR versus prediabetes-IR *(χ^2^* = 14.780, df = 7, p = 0.031, Nagelkerke = 0.364) with an accuracy of 71.2%, sensitivity of 74.4%, and specificity of 72.7%.

**Table 4:** Results of two different binary logistic regression analyses with Prediabetes (versus healthy controls) and Prediabetes-IR versus Prediabetes+IR as dependent variables and the biomarkers as explanatory variables.

| Dichotomies | Explanatory variables | B | SE | Wald | df | p | OR | 95% CI |
| --- | --- | --- | --- | --- | --- | --- | --- | --- |
| **Prediabetes / Controls** | IL-10 | 0.359 | 0.128 | 7.876 | 1 | 0.005 | 1.431 | 1.114-1.839 |
| | zLnβEP | 0.316 | 0.137 | 5.311 | 1 | 0.021 | 1.372 | 1.048-1.795 |
|  | zLnEM2 | 0.846 | 0.396 | 4.569 | 1 | 0.033 | 2.331 | 1.073-5.063 |
| **Prediabetes-IR / Prediabetes+IR** | IL-10 | 0.618 | 0.05 | 2.505 | 1 | 0.043 | 1.083 | 0.981-1.195 |
|  | zLnEM2/KOR | 0.555 | 0.433 | 4.846 | 1 | 0.024 | 2.611 | 1.187-3.947 |

zLn: z-score of the natural logarithm, IL: interleukin; KOR: κ-opioid receptor; MOR: $\mu$ opioid receptor; EM2: Endomorphin-2; βEP: β-endorphin; IR: insulin resistance (HOMA2IR>2.5)

## Correlation

### Prediction of Symptom Domains by Biomarkers

Table 5 shows different stepwise multiple regression analyses with the IR parameters as dependent variables and the eight biomarkers as explanatory variables while allowing the effects of education, age, and sex. Regression #1 showed that 21.1% of the variance in the total FPG can be explained by the regression on IL-10, zLnKOR, and zLn (EM2/KOR). Regressions #2, #4, #5, and #6 showed that the same variables explained a considerable part of the variance in insulin (22.4%), HbA1c (29.0%), HOMA2IR (29.3%), and HOMA2%S (29.7%). Regression #3 showed that 13.0% of the variance in *I/G* ratio was explained by IL-10. Regression #7 showed that zLnHOMA%B cannot be explained by any measured biomarker (p = 0.742).

**Table 5:**
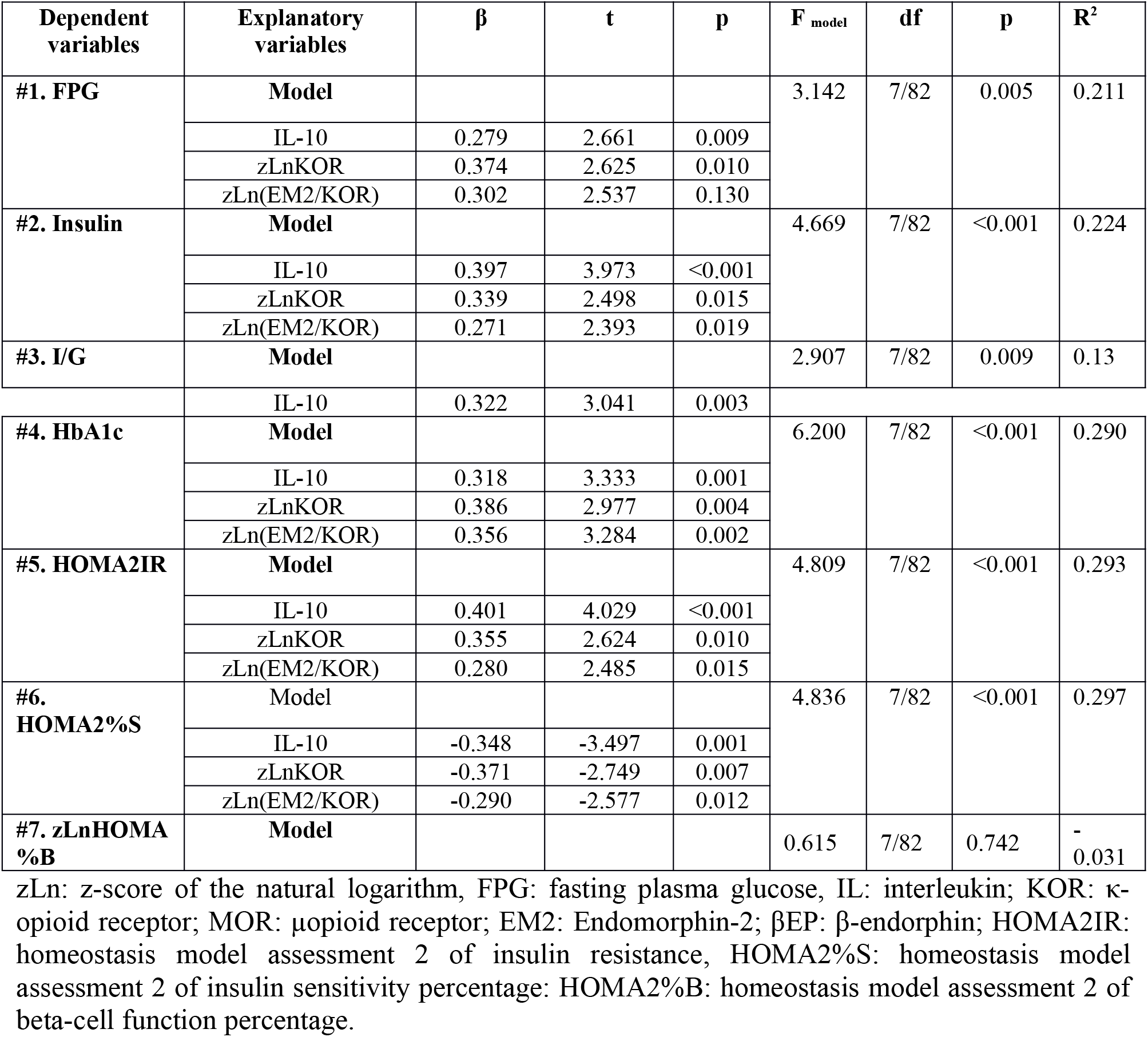
Results of multiple regression analysis with IR parameters as dependent variables and biomarkers as explanatory variables.

## Discussion

This case–control observational study was designed to investigate the possible role of opoiod system in the pathogenesis of prediabetes. Accordingly, this problem was solved in this study. Dyslipidemia and increased atherogenic indices were observed in the prediabetes+IR group compared with the controls, indicating the negative effect of elevated IR on the lipid metabolism (Table 1). IR can also alter systemic lipid metabolism, leading to the generation of dyslipidemia. Along with endothelial dysfunction, which can also be created by an abnormality of insulin signaling, IR contributes to the atherosclerotic plaque production [36]. IR in the myocardium produces damage by signal transduction modification, impaired regulation of substrate metabolism, and variation in the delivery of substrates to the myocardium [36]. These variations are engaged in the presence of increased IR in patients with prediabetes, indicating the huge effect of the elevated IR on the lipid metabolism.

The first main finding is the elevation in the levels of the endogenous opioid and their receptors in the prediabetes groups compared with the controls. These results signify the dependence of these parameters on the glucose metabolism state more than its dependence on the response to insulin hormone. To our knowledge, this study was the first to deal with the endogenous opioid in prediabetes subjects. Some previous studies have dealt with the opioid peptides and receptors in the IR state in animals [28, 37] and humans [38, 39]. MOR is enhanced on insulin-sensitive tissues, such as skeletal muscle, originating in an inversion of the impairment of insulin-stimulated glucose disposal in genetically obese Zucker rats via exercise training [37]. This enhancement of IR is associated with the increased circulation of ß-endorphin, thus enhancing the post receptor and insulin signaling cascade, including the downstream effectors of the phosphatidylinositol 3-kinase (PI3-kinase) signaling pathway implicated in the translocation of the glucose transporter [28, 37]. The increase in βEP release from the adrenal gland may stimulate peripheral opioid MOR to boost the expression of muscle glucose transporters and/or reduce hepatic gluconeogenesis at the gene level, thereby directing to enhance glucose utilization in peripheral tissues for the improvement of severe hyperglycemia [28].

Insulin sensitivity is improved through the peripheral MOR stimulation, overcoming defects in insulin signal transduction associated with the post receptor in IRS-1 associated with the PI3-kinase step. Insulin sensitivity is enhanced through the peripheral MOR activation. Opioid receptor stimulation, especially that of the μ-subtype, may supply merits in the improvement of defective insulin function. The opioid system manages the endocrine function of β-cells [40]. Within the pancreas, EOPs result in insulin secretion via the paracrine and the intracrine mechanisms [41]. Opioid activity can perform an important role in the physiopathology of hyperinsulinemia in hyperandrogenic women [38]. Hence, locally increased opioid activity results in insulin release from the pancreas. IR is a consequence of the downregulation of hepatic insulin receptors, thereby increasing the insulin levels in peripheral blood [6, 42] Sinaiko and Caprio 2012). Interestingly, brain damage and white matter disruption occur in the IR stage without overt diabetes, and these structural abnormalities may be involved in early cognitive impairment [43] and secrete endogenous peptides to the circulation and increase their levels. Patients with insulin-resistant conditions (like polycystic ovarian syndrome) have greater limbic MOR availability than the controls. Moreover, receptor availability is associated with the severity of IR, and MOR availability is normalized after insulin-regulating therapy [39].

The typical brain areas for opioid neurotransmission are the amygdala and nucleus accumbens, which affect the management of appetite and mood.

The elevation of EM2 in prediabetes subjects may be due to the increase in their release in response to any possible hurt of the beta cells. In animal studies, EM2 can maintain the betacell islets from the injury caused by streptozotocin, alloxan, and hydrogen peroxide [44]. EM2 enhances the viability of islet and elevates the insulin accumulation of the cell supernatant after streptozotocin and alloxan stimulation. Our observations imply that endomorphins may have protective effects on islet cell oxidative injury [44]. When the μ and κ opioid antagonists are microinjected into the hypothalamic paraventricular nucleus or the nucleus accumbens of rats, they decrease food intake under deprivation states. MOR and KOR seem accountable for the inclusion of various food intake by the nucleus [45]. The outcomes also propose that MOR and KOR may be altered by blood glucose levels, possibly participating cellular energetics-mediated variation in potassium channels in female rats [46]. Moreover, elevated insulin levels lead to the variation in the sensitivity of KOR and MOR to nociception by agonists [47]. The blood glucose of diabetic mice decreases significantly behind the therapy with a selective KOR agonist. This phenomenon is time-dependent and related to the corresponding variation of GLUT4 translocation in the skeleton muscles. Hence, the enhancement of KOR decreases the hyperglycemia in streptozotocin-induced diabetic mice [48]. Insulin elevates the efficacy of KOR motivation by phosphorylating two tyrosine residues in the first and second intracellular loops of the receptor. Thus, tyrosine phosphorylation may supply an influential mechanism for the variation of G protein-coupled receptor signaling [49]. Together, these studies support that the insulin in the CNS may reduce the effect of CNS KOR system(s) that mediate palatable feeding [50]. Our results are in accordance with the mutual effect of IR and hyperinsulinemia with the endogenous opioid receptors. The presence of these changes in the prediabetic groups in comparison with controls should be considered when seeking for a new treatment target to prevent the development of prediabetes into T2DM.

Serum IL-10 was significantly different among the three study groups, and the score followed the order: controls < prediabetes-IR < prediabetes+IR. In a previous study, the serum IL-10 level is elevated in T2DM but not in prediabetes compared with the healthy controls [51]. However, this study is not choosing a homogenized group in number, and patients with positive CRP were excluded. The decreased levels of the proinflammatory cytokine IL-12 in the prediabetes stage and the increased levels of the anti-inflammatory cytokine IL-10 in the T2DM stage suggest a continuous organismal effort for halting the activation of the Th1 subpopulation that acts as a main generator of proinflammatory INFγ, possibly as a mechanism to maintain homeostasis in relevant tissues. Notably, several studies have reported the addition of the Th1 cell subset and causal participation of this phenomenon in inflammation and IR in the mouse models of diabetes [52]. IL-6 was significantly elevated in the prediabetes groups compared with the control group, whereas no such difference was observed between the prediabetes subgroups. IL-6 has exhibited no significant variation between the prediabetes groups and controls [53]. However, increased IL-6, another proinflammatory marker, is observed in hyperglycemic/hyperinsulinemic conditions [54, 55]. The studies on inflammatory markers have indicated that participants with a combined increase of IL-6 and IL-1β have approximately threefold elevated risk of developing diabetes, whereas low levels of IL-1β alone exhibit no substantial elevation in risk [56].

Results exhibited significant associations between IL-10 and MOR and between IL-6 and KOR. The connection among endogenous opioids and immunosuppression has been explored in vitro and in vivo. However, results are conflicting. Exogenous narcotics hinder the capacity of macrophages, natural killer cells, and T-cells to weaken the gut barrier in vitro and in animal models [57]. High dosage and the commencement of opioid therapy for pain correspond with a high danger of infectious diseases, such as pneumonia. However, immune cells secrete endogenous opioid peptides, which are linked to peripheral opioid receptors to relieve inflammatory pain. In general, the immune framework and endogenous opioids have an equal collaboration (57). IL-10 and MOR has a relationship in depression [58], indicating that the mood causes on such association in addition to the suggested correlation between immune and opioid systems.

The second major finding was obtained from the multivariate GLM analysis, which removed the effects of the covariates (age, BMI, sex, education, IR parameters, physical activity, and smoking) to compare the parameters in accordance with the type of diagnosis (Table 2). Approximately 20% (0.198) of the values of immune–opioid biomarkers can be explained by the presence of prediabetes and IR. Based on the results on Tables 2 and 3 (SE), all eight biomarkers were significantly higher in patients with prediabetes compared with the controls. Among all the examined biomarkers, zLnβEP had the highest effect on the diagnosis of prediabetes. βEP cells have been demonstrated in areas surrounding the pancreatic β-cells, and opioids enhance insulin secretion [42]. The improvement of IR is associated with the increased circulating β-endorphin to ameliorate the postreceptor insulin signaling cascade and the improvement in insulin sensitivity through the peripheral MOR motivation [59]. MOR manages body weight by sharing insulin release, exhibiting a novel target for new diabetes treatments [60].

The endogenous opioid system negatively manages insulin secretion from isolated Langerhans islets in vitro [40]. In vivo, the β-endorphin also weakens insulin secretion when administered through intravenous infusion [61]. In addition, some groups have reported the dual stimulatory/inhibitory action of β-endorphin on insulin secretion depending on dose, obesity condition, or circulating glucose level [62]. MORs act as a part of the complex opioid system and mediate the effects of endogenous opioids, such as endomorphin-2 and various exogenous opioid agonists [63]. The determination of the endogenous opioid peptide expression in the Langerhans islet and peptide secretion after glucose motivation [64]. transform the endocrine pancreas as a potential candidate for MOR management. MOR participates in glucose homeostasis by negatively regulating glucose tolerance by inhibiting insulin release from the β-cell and decreasing β-cell mass [60].

The binary logistic regression analyses in Table 4 indicated that the increased levels of the combination of IL-10, zLnβEP, and zLnEM2 can be used to differentiate prediabetes+IR from other groups with good accuracy (76.7%). Another compilation of increased IL-10 and zLnEM2/KOR can be used to predict prediabetes+IR with good accuracy (71.2%). These results further indicate the importance of these parameters in prediabetes+IR and the strong correlation with the immune-opioid system. Endogenous opioid activity increases by acting on the pancreatic B cells, an increase in insulin secretion, resulting in elevated insulin levels in the hepatic circulation. Elevated insulin levels lead to a downregulation of hepatic insulin receptors, thereby causing and maintaining hyperinsulinemia [65]. Morbid obesity is related to reduced MOR availability in the brain, and the endogenous opioid system is a key component underlying human obesity [66]. The increase in body mass is associated negatively with MOR availability [66]. The central stimulation of the MOR impairs glucose tolerance and insulin sensitivity and motivates hepatic gluconeogenesis [67]. The stimulation of peripheral MOR can enhance IR in animals and provide a novel target for the treatment of IR [68]. MOR activation may ameliorate IL-6-induced IR through various insulin signals opposite the responsiveness of insulin [69].

Mice lacking MOR in the whole body exhibit increased glucose tolerance and insulin sensitivity and elevated fatty acid oxidation when fed with a high-fat diet [67]. The stimulation of KOR results in the reduction in IL-6 levels [70]. Table 5 shows an important result on the effect of the measured biomarkers on the IR parameters. Approximately one-fifth (21.1%) of the variance in FPG can be explained by the regression on IL-10, zLnKOR, and zLn(EM2/KOR). The elevated endogenous opiates may affect the insulin response to glucose in subjects with obesity through impaired or normal oral glucose tolerance test [71]. Same changes explain an important part of the variations in insulin, HbA1c, HOMA2IR, and HOMA2%S. Opioid activity can play a critical role in the physiopathology of hyperinsulinemia in hyperandrogenic women [38]. The βEP level is not affected during the oral glucose tolerance test. Furthermore, reducing the IR condition after the reduction in body weight has not affected the βEP level [72]. In healthy persons, the independent parameters (FBG, insulin, and HbA1c) associated well because the beta cells worked properly. Elevated blood glucose stimulates the elevation in insulin concentrations as released from the healthy beta cells to keep normal blood glucose concentrations over long periods [73]. The hyperproduction of endogenous opioid peptides in obesity increases insulin release motivated by food intake but does not appreciably affect the insulin release stimulated by circulating glucose or amino acids [74].

## Conclusion

The subjects with prediabetes had dyslipidemia, but all of them underwent the IR state. IL-10, βEP, MOR, and EM2 were significantly higher in the prediabetes subgroups than in the control group. Among all the examined biomarkers, βEP had the highest effect on the diagnosis of prediabetes. Serum IL-10 increased among the three study groups, and the score followed the order: controls < prediabetes-IR < prediabetes+IR. Serum IL-6 was significantly increased in the prediabetes groups compared with the control group, and the increase was independent of the IR. MOR was correlated with IL-10 and KOR, and KOR was correlated with IL-6 and EM2. Prediabetes+IR can be predicted using the increased levels of the combination of IL-10, βEP, and EM2 and by a combination of IL-10 and EM2/KOR with good sensitivity and specificity. A significant percentage of FPG, insulin, HbA1c, HOMA2IR, and HOMA2%S can be explained by the regression on IL-10, KOR, and EM2/KOR in the prediabetes group. Overall, the opioid peptides and their receptors were upregulated in prediabetes depending on the significance of IR. These changes in the opioid system are dependent on the immune cytokines.

## Limitations of the Study

The results of this study should be discussed with respect to its limitations. First, the delta opioid receptor, T regulatory phenotypes, and some other cytokines, including those produced by M1 macrophage, T helper cells 1, 2, and 17 should be measured to examine the associations of opioids with the full spectrum of cytokines. Second, the effects of dynorphin and β-endorphin on the production of cytokines and the effects of immune activation (by mitogen or LPS stimulation) on KOR–MOR protein production should be examined by stimulated and unstimulated whole blood cultures of patients and controls.

## Abbreviations

(FBG): fasting blood glucose
(IR): insulin resistance
(ELISA): Enzyme linked immmunosorbent assay
(IL): interleukin
(MOR and KOR): μ- and κ-opioid receptors
(EM2): endomorphin-2
(βEP): β-endorphin

## Acknowledgement

We thank all the study volunteers for their contribution.

## References

1 Buysschaert M, Bergman M. Definition of prediabetes. Med Clin North Am. 2011; 95(2):289–297.

2 YanFeng Li, Linda S. Geiss, Nilka R. Burrows, Deborah B. Rolka, Ann Albright, Div of Diabetes Translation, et al. Awareness of Prediabetes - United States, 2005-2010. MMWR Morb Mortal Wkly Rep. 2013; 62(11):209–212.

3 Centers for Disease Control and Prevention, U.S. Department of Health and Human Services; 2017. National Diabetes Statistics Report 2017; 2017 (cited 2020 Jun 2). Available from: https://www.cdc.gov/diabetes/data/statistics-report/index.html.

4 American Diabetes Association. Classification and Diagnosis of Diabetes. Diabetes Care. 2017; 40:(S11-s24).

5 Dall T, Thiselton D, Varvel S. Targeting insulin resistance: the ongoing paradigm shift in diabetes prevention. Am J Managed Care. 2013; 19(2):E7.

6 Sinaiko AR, Caprio S. Insulin Resistance. J Pediatr. 2012; 161(1):11–15.

7 Khetan AK, Rajagopalan S. Prediabetes. Can J Cardiol. 2018; 34(5):615–623.

8 Singh Y, Garg MK, Tandon N, Marwaha RK. A study of insulin resistance by HOMA-IR and its cut-off value to identify metabolic syndrome in urban Indian adolescents. J Clin Res Pediatr Endocrinol. 2013; 5(4):245–251.

9 Ziaee A, Esmailzadehha N, Oveisi S, Ghorbani A, Ghane L. The threshold value of homeostasis model assessment for insulin resistance in Qazvin Metabolic Diseases Study (QMDS): assessment of metabolic syndrome. J Res Health Sci. 2015; 15(2):94–100.

10 Al-Hakeim HK, Abdulzahra MS. Correlation Between Glycated Hemoglobin and Homa Indices in Type 2 Diabetes Mellitus: Prediction of Beta-Cell Function from Glycated Hemoglobin. J Med Biochem. 2015; 34(2):191–199.

11 Chua LM, Lim ML, Chong PR, Hu ZP, Cheung NS, Wong BS. Impaired neuronal insulin signaling precedes Aβ42 accumulation in female AβPPsw/PS1ΔE9 mice. JAlzheimer’s Dis. 2012; 29(4):783–791.

12 Macklin L, Griffith CM, Cai Y, Rose GM, Yan XX, Patrylo PR. Glucose tolerance and insulin sensitivity are impaired in APP/PS1 transgenic mice prior to amyloid plaque pathogenesis and cognitive decline. Exp Gerontol. 2017; 88:9–18.

13 de Felice FG. Alzheimer’s disease and insulin resistance: Translating basic science into clinical applications. J Clin Investig. 2013; 123(2):531–539.

14 Bomfim TR, Forny-Germano L, Sathler L B, Brito-Moreira J, Houzel JC, Decker H, et al. An anti-diabetes agent protects the mouse brain from defective insulin signaling caused by Alzheimer’s disease-associated A_oligomers. JClin. Investig. 2012; 22: 1339–1353.

15 Reijmer YD, Leemans A, Brundel M, Kappelle LJ, Biessels GJ, Utrecht. Vascular Cognitive Impairment Study Group. Disruption of the cerebral white matter network is related to slowing of information processing speed in patients with type 2 diabetes. Diabetes. 2013; 62(6):2112–2115.

16 Ekblad LL, Rinne JO, Puukka PJ, Laine HK, Ahtiluoto SE, Sulkava RO, et al. Insulin resistance is associated with poorer verbal fluency performance in women. Diabetologia. 2015; 58(11):2545–2553.

17 Reid RL, Yen SS. beta-Endorphin stimulates the secretion of insulin and glucagon in humans. J Clin Endocrinol Metab. 1981; 52(3):592–594.

18 Ipp E, Dobbs R, Unger RIt. Morphine and beta-endorphin influence the secretion of the endocrine pancreas. Nature. 1978; 276(5684):190–191.

19 Giugliano D, Ceriello A, Salvatore T, Paolisso G, D’Onofrio F, Lefèbvre P. Beta-endorphin infusion restores acute insulin responses to glucose in type-2 diabetes mellitus. J Clin Endocrinol Metab. 1987; 64(5):944–948.

20 Zhang M, Zheng M, Schleicher RL. Localization of b-endorphin in rabbit pancreatic islets. Mol Cell Neurosci. 1992; 3(6):536–547.

21 Nakao K, Nakai Y, Jingami H, Oki S, Oki S, Fukata J, Imura H. Substantial rise of plasma b-endorphin levels after insulin-induced hypoglycemia in human subjects. J Clin Endocrinol Metab. 1979; 49(6):838–841.

22 Hickey MS, Trappe SW, Blostein AC, Edwards BA, Goodpaster B, Craig BW. et al. Opioid antagonism alters blood glucose homeostasis during exercise in humans. J Appl Physiol. 1994; 76(6):2452e60.

23 Chen X, Wang L, Fan S, Song S, Min H, Wu Y, et al. Puerarin acts on the skeletal muscle to improve insulin sensitivity in diabetic rats involving μ-opioid receptor. Eur J Pharmacol. 2018; 818:115–123.

24 Givens JR, Wiedemann E, Andersen RN, Kitabchi AE. beta-Endorphin and beta-lipotropin plasma levels in hirsute women: correlation with body weight. J Clin Endocrinol Metab. 1980; 50(5):975–976.

25 Curran AM, Scott-Boyer MP, Kaput J, Ryan MF, Drummond E, Eileen R Gibney ER, et al. A proteomic signature that reflects pancreatic beta-cell function. PLoS One 2018; 13(8):e0202727.

26 Wen T, Peng B, Pintar JE. The MOR-1 opioid receptor regulates glucose homeostasis by modulating insulin secretion. Mol Endocrinol. 2009; 23(5):671e8.

27 Najafipour H, Beik A. The impact of opium consumption on blood glucose, serum lipids and blood pressure, and related mechanisms. Front Physiol. 2016 13;7:436. eCollection 2016.

28 Liu IM, Cheng JT. Mediation of endogenous β-endorphin in the plasma glucose-lowering action of herbal products observed in type1-like diabetic rats. Evid Based Complement Alternat Med. 2011; 2011:987876.

29 Vuong C, Van Uum SH, O’Dell LE, Lutfy K, Friedman TC. The effects of opioids and opioid analogs on animal and human endocrine systems. Endocr Rev. 2010; 31(1):98–132.

30 Song CY, Bi HM. Effects of puerarin on plasma membrane GLUT4 content in skeletal muscle from insulin-resistant Sprague-Dawley rats under insulin stimulation. Zhongguo Zhong Yao Za Zhi. 2004; 29(2):172–175.

31 Wang CL, Diao YX, Xiang Q Ren YK3, Gu N. Diabetes attenuates the inhibitory effects of endomorphin-2, but not endomorphin-1 on gastrointestinal transit in mice. Eur J Pharmacol. 2014; 738:1–7.

32 Kurauti MA, Costa-Júnior JM, Ferreira SM, Santos GJ, Sponton CHG, Carneiro EM, et al. Interleukin-6 increases the expression and activity of insulin-degrading enzyme. Sci Rep. 2017; 7:46750.

33 Rehman K, Akash MSH, Liaqat A, Kamal S, Qadir MI, Rasul A. Role of Interleukin-6 in Development of Insulin Resistance and Type 2 Diabetes Mellitus. Crit Rev Eukaryot Gene Expr. 2017; 27(3):229–236.

34 de Waal Malefyt R, Abrams J, Bennett B, Figdor CG, de Vries JE. Interleukin 10(IL-10) inhibits cytokine synthesis by human monocytes: an autoregulatory role of IL-10 produced by monocytes. J Exp Med. 1991; 174(5):1209–1220.

35 Leon-Cabrera S, Arana-Lechuga Y, Esqueda-León E et al. Reduced systemic levels of IL-10 are associated with the severity of obstructive sleep apnea and insulin resistance in morbidly obese humans. Mediators Inflamm. 2015; 2015:493409.

36 Ormazabal V, Nair S, Elfeky O, Aguayo C, Salomon C, Zuñiga FA. Association between insulin resistance and the development of cardiovascular disease. Cardiovasc Diabetol. 2018; 17(1):122.

37 Cheng KC, Asakawa A, Li Y, Liu IM, Amitani H, Cheng JT, Inui A, Opioid μ-receptors as new target for insulin resistance. Pharmacol Ther. 2013; 139(3):334–340.

38 Sir T, López G, Alba F, et al. Effect of chronic blockade of the opiodergic receptor on insulin resistance in a hyperandrogenic woman. Rev Med Chil. 1994; 122(4):441–447.

39 Berent-Spillson A, Love T, Pop-Busui R et al. Insulin resistance influences central opioid activity in polycystic ovary syndrome. Fertil Steril. 2011; 95(8):2494–2498.

40 García-Barrado MJ, Iglesias-Osma MC, Rodríguez R, Martín M, Moratinos J. Role of μ-opioid receptors in insulin release in the presence of inhibitory and excitatory secretagogues. Eur J Pharmacol. 2002; 448(1):95–104.

41 Bottcher B, Seeber B, Leyendecker G, Wildt L. Impact of the opioid system on the reproductive axis. Fertility and Sterility. 2017; 108(2):207–213.

42 Hadziomerovic D, Rabenbauer B, Wildt L. Normalization of hyperinsulinemia by chronic opioid receptor blockade in hyperandrogenemic women. Fertil Steril. 2006; 86(3), 651–657.

43 Willette A.A, Xu G, Johnson SC et al. Insulin resistance, brain atrophy, and cognitive performance in late middle-aged adults. Diabetes Care. 2013; 36(2):443–449.

44 Tian LM, Liu J, Sun XL, Gao CX, Fan Y, Guo Q. A protective effect of endomorphins on the oxidative injury of islet. Exp Clin Endocrinol Diabetes. 2010; 118(8):467–472.

45 Ragnauth A, Ruegg H, Bodnar RJ. Evaluation of opioid receptor subtype antagonist effects in the ventral tegmental area upon food intake under deprivation, glucoprivic and palatable conditions. Brain Res. 1997; 767(1):8–16.

46 Tandon M, Srivastava RK, Nagpal RK, Khosla P, Singh J. Differential modulation of nociceptive responses to mu and kappa opioid receptor directed drugs by blood glucose in experimentally induced diabetes rats. Indian J Exp Biol. 2000; 38(3):242–248.

47 Srivastava RK, Verma S, Tandon M. Effect of insulin hypoglycemic stress on nociceptive responses to mu- and kappa-opioid receptor agonists at LH-surge in female rats. Methods Find Exp Clin Pharmacol. 2004; 26(3):189–194.

48 Shang Y, Guo F, Li J, et al. Activation of κ-opioid receptor exerts the glucose-homeostatic effect in streptozotocin-induced diabetic mice. J Cell Biochem, 2015; 116(2):252–259.

49 Appleyard SM, McLaughlin JP, Chavkin C. Tyrosine phosphorylation of the kappa-opioid receptor regulates agonist efficacy. J Biol Chem. 2000; 275(49):38281–38285.

50 Sipols AJ, Bayer J, Bennett R, Figlewicz DP. Intraventricular insulin decreases kappa opioid-mediated sucrose intake in rats. Peptides. 2002; 23(12):2181–2187.

51 Nunez Lopez YO, Garufi G, Seyhan AA. Altered levels of circulating cytokines and microRNAs in lean and obese individuals with prediabetes and type 2 diabetes. Mol Biosyst. 2016; 13(1):106–121.

52 Kornete M, Mason ES, Piccirillo CA. Immune Regulation in T1D and T2D: Prospective Role of Foxp3+Treg Cells in Disease Pathogenesis and Treatment. Front Endocrinol(Lausanne). 2013; 4:76.

53 Butkowski EG, Jelinek HF. Hyperglycaemia, oxidative stress and inflammatory markers. Redox Rep. 2017; 22(6):257–264.

54 Fishel MA, Watson G, Montine TJ et al. Hyperinsulinemia provokes synchronous increases in central inflammation and β-amyloid in normal adults. Arch Neurol. 2005; 62(10):1539–1544.

55 Butkowski EG, Brix LM, Al-Aubaidy H, Kiat H, Jelinek HJ. Diabetes, oxidative stress and cardiovascular risk. J Med Clin Sci. 2016; 5(1):17–23.

56 Spranger J, Kroke A, Möhlig M et al. Inflammatory cytokines and the risk to develop type 2 diabetes: results of the prospective population-based European Prospective Investigation into Cancer and Nutrition (EPIC)-Potsdam study. Diabetes. 2003; 52(3):812–817.

57 Plein LM, Rittner HL. Opioids and the immune system - friend or foe. Br J Pharmacol. 2018; 175(14):2717–2725.

58 Al-Fadhel SZ, Al-Hakeim HK, Al-Dujaili AH, Maes M. IL-10 is associated with increased mu-opioid receptor levels in major depressive disorder. Eur Psychiatry. 2019; 57:46–51.

59 Kai-Chun Cheng, Akihiro Asakawa, Ying-Xiao Li et al. Opioid μ-receptors as new target for insulin resistance. Pharmacol Ther. 2013; 139(3):334–340.

60 Ting Wen, Bonnie Peng, John E. Pintar. The MOR-1 Opioid Receptor Regulates Glucose Homeostasis by Modulating Insulin Secretion. Mol Endocrinol. 2009; 23(5):671–678.

61 Giugliano D, Cozzolino D, Ceriello A, Salvatore T, Paolisso G, Torella R. β-Endorphin and islet hormone release in humans: evidence for interference with cAMP. Am J Physiol. 1989; 257(3 Pt 1):E361–E366.

62 Khawaja XZ, Green IC. Dual action of β-endorphin on insulin release in genetically obese and lean mice. Peptides. 1991; 12(2):227–233.

63 Henriksen G, Willoch F. Imaging of opioid receptors in the central nervous system. Brain. 2008; 131(Pt 5):1171–1196.

64 Josefsen K, Buschard K, Sorensen LR, Wøllike M, Ekman R, Birkenbach M 1998. Glucose stimulation of pancreatic β-cell lines induces expression and secretion of dynorphin. Endocrinology. 1998; 139(10):4329–4336.

65 Mahabeer S, Jialal I, Norman RJ, Naidoo C, Reddi K, Joubert SM. Insulin and C-peptide secretion in non-obese patients with polycystic ovarian disease. Horm Metab Res. 1989; 21(9):502–506.

66 Henry K. Karlsson, Lauri Tuominen, Jetro J. Tuulari et al. Obesity Is Associated with Decreased MOR-Opioid But Unaltered Dopamine D2 Receptor Availability in the Brain. J Neurosci. 2015; 35(9): 3959–3965.

67 Tudurí E, Nogueiras R. Mu opioid receptor: from pain to glucose metabolism. Oncotarget. 2017; 8(4):5643–5644.

68 Tzeng TF, Lo CY, Cheng JT, Liu IM. Activation of mu-opioid receptors improves insulin sensitivity in obese Zucker rats. Life Sci. 2007; 80(16):1508–1516.

69 Tzeng TF, Liu IM, Cheng JT. Activation of opioid mu-receptors by loperamide to improve interleukin-6-induced inhibition of insulin signals in myoblast C2C12 cells. Diabetologia. 2005; 48(7):1386–1392.

70 Zhou X, Wang D, Zhang Y, Zhang J, Xiang D, Wang H. Activation of κ-opioid receptor by U50,488H improves vascular dysfunction in streptozotocin-induced diabetic rats. BMC Endocr Disord. 2015; 28: 15:7.

71 Satta MA, Maussier ML, Scoppola A, Menini E, Liberale I, Lio S, Monaco F, Roche J. Endogenous opiate modulators of insulin secretion in the obese. C R Seances Soc Biol Fil. 1988; 182(6):538–543.

72 Ritter MM, Sönnichsen AC, Möhrle W, Richter WO, Schwandt P. Beta-endorphin plasma levels and their dependence on gender during an enteral glucose load in lean subjects as well as in obese patients before and after weight reduction. Int J Obes. 1991; 15(6):421–427.

73 Hellman B, Grapengiesser E. Glucose-induced inhibition of insulin secretion. Acta Physiol (Oxf). 2014; 210(3):479–88.

74 Vettor R, Martini C, Cestaro S, Manno M, Sicolo N, Federspil G. Possible involvement of endogenous opioids in beta-cell hyperresponsiveness in human obesity. Int J Obes. 1989; 13(4):425–432.

